# Effect of knee joint loading on chondrocyte mechano-vulnerability and severity of post-traumatic osteoarthritis induced by ACL-injury in mice

**DOI:** 10.1101/2021.06.16.448294

**Authors:** Alexander Kotelsky, Anissa Elahi, Nejat Yigit Can, Ashley Proctor, Sandeep Mannava, Christoph Pröschel, Whasil Lee

**Affiliations:** Department of Biomedical Engineering, University of Rochester, Rochester, NY 14620, USA; Department of Pharmacology and Physiology, University of Rochester Medical Center, Rochester, NY 14620, USA; Center for Musculoskeletal Research, University of Rochester Medical Center, Rochester, NY 14620, USA; Department for Biomedical Genetics, University of Rochester, Rochester, NY 14620, USA; Department of Orthopaedics, University of Rochester Medical Center, Rochester, NY 14620, USA

**Author notes:** Address correspondence to: W. Lee, Department of Biomedical Engineering & Pharmacology and Physiology, University of Rochester;, 601 Elmwood Ave. Box 711 Rm 4.8553Rochester, NY14620, USA.

**Keywords:** Post-traumatic Osteoarthritis (PT-OA), anterior cruciate ligament (ACL)-injury, chondrocyte death, extracellular matrix (ECM), gait analysis

## Abstract

**Objective:** The objective of this study is to understand the role of altered *in vivo* mechanical environments in knee joints post anterior cruciate ligament (ACL)-injury in chondrocyte vulnerability against mechanical stimuli and in the progression of post-traumatic osteoarthritis (PT-OA).

**Methods:** Differential *in vivo* mechanical environments were induced by unilateral ACL-injury (uni-ACL-I) and bilateral ACL-injury (bi-ACL-I) in 8-week-old female C57BL/6 mice. The gait parameters, the mechano-vulnerability of *in situ* chondrocytes, Young’s moduli of cartilage extracellular matrix (ECM), and the histological assessment of OA severity (OARSI score) were compared between control and experimental groups at 0∼8-weeks post-ACL-injury.

**Results:** We found that bi-ACL-I mice experience higher joint-loading on their both injured limbs, but uni-ACL-I mice balance their joint-loading between injured and uninjured hind limbs resulting in a reduced joint-loading during gait. We also found that at 4- and 8-week post-injury the higher weight-bearing hind limbs (i.e., bi-ACL-I) had the increased area of chondrocyte death induced by impact loading and higher OARSI score than the lower weight-bearing limbs (uni-ACL-I). Additionally, we found that at 8-weeks post-injury the ECM became stiffer in bi-ACL-I joints and softer in uni-ACL-I joints.

**Conclusions:** Our results show that ACL-injured limbs with lower *in vivo* joint-loading develops PT-OA significantly slower than injured limbs with higher joint-loading during gait. Our data also indicate that articular chondrocytes in severe PT-OA are more fragile from mechanical impacts than chondrocytes in healthy or mild PT-OA. Thus, preserving physiologic joint-loads on injured joints will reduce chondrocyte death post-injury and may delay PT-OA progression.

## Introduction

Joint injury is a major risk factor of symptomatic osteoarthritis (OA), a prevalent and debilitating condition of load-bearing joints characterized by progressive degeneration of cartilage extracellular matrix (ECM)^1-7^. Anterior cruciate ligament (ACL)-tear is the most common knee injury and more than 50% of ACL-injury patients develop post-traumatic (PT)-OA within 5∼20 years post-injury regardless of whether or not they have ACL-reconstruction surgery^8-12^. Unfortunately, the majority of injuries occur in young adults between the ages of 15 and 24^13-17^, and these young adults with ACL injuries are more likely to develop OA before the age of 40^11, 18^. Risk of ACL-revision surgery is especially pronounced in younger patients, who are subject to multiple revision surgeries over a lifetime, and who may have only options of surgical total knee joint replacement to treat PT-OA^19, 20^. The hallmark of ACL-injury is knee joint destabilization. ACL, a ligament directly connecting the femur to the tibia, stabilizes the knee joint in the anterior-to-posterior direction, prevents anterior-tibial subluxation, as well as provides rotational stability. A sudden turn or non-contact mechanism typically causes an ACL tear resulting in knee subluxation, pivot shift, and joint instability indicated in altered joint kinematics and gait patterns^21-27^.

Mechanical factors heavily influence chondrocyte’ biosynthetic activities and ECM homeostasis, and chondrocyte mechanotransduction plays a critical role in the pathogenesis of OA^6, 28-33^. Thus, the abnormal mechanical loading is presumed to contribute to OA progression post-ACL-injury. In addition, ACL-injuries often lead to more severe OA progression due to the concomitant damages on the meniscus, other ligaments, or articular cartilage^34-37^. Furthermore, contralateral non-injured knee joints have high risks of joint injuries due to the compensatory aberrant joint loading^38, 39^, and 12% of ACL-injured patients have recurring ACL ruptures in their contralateral knees within 2 to 5 years post-injury^40, 41^. These combinations of damages may cause individuals to load different degrees of abnormal mechanical stimuli on their ACL-injured joints, thus may lead to diverse rates of PT-OA progression.

Several murine PT-OA studies have identified the increased chondrocyte death in injured cartilage by histological assessment^42-44^. Since promoting chondrocyte survival after joint injury may delay cartilage degradation^45-49^, it is important to understand how the abnormal *in vivo* mechanical loads alter chondrocyte viability and cartilage homeostasis post-injuries. In addition, the ability of chondrocytes to withstand injurious forces, termed ‘mechano-vulnerability’, depends on mechano-sensitivity of chondrocytes and mechanical properties of the ECM. Currently, a limited information is known about the effects of *in vivo* loading on chondrocyte viability post-injury, and loading-dependent chondrocyte mechano-vulnerability and the mechanical properties of ECM have not been studied over the PT-OA progression post-ACL-I.

Here, we investigate the extent to which alterations in *in-vivo* mechanical loading environments impact chondrocyte mechano-vulnerability in ACL-injured knees using a non-invasive ACL injury mouse model. The alterations in *in vivo* mechanical environment were induced through unilateral (uni-ACL-I) and bi-lateral (bi-ACL-I) ACL injuries of hind legs. We found that uni-ACL-I mice experienced lower weight-bearing in their hind paws during the gait as compared to bi-ACL-I mice. We also found that articular cartilage of ACL-injured mice with higher weight-bearing (i.e., bi-ACL-I) had more mechanically vulnerable chondrocytes and more severe cartilage degradation, the evidence of PT-OA, as compared to injured mice with lower gait-associated weight-bearing (i.e., uni-ACL-I). Our findings demonstrate that it is critical to avoid abnormal joint mechanics and reduce chondrocyte mechano-sensitivity post-ACL-I as a potential treatment aimed to delay or halt PT-OA progression.

## Methods

### Non-invasive ACL injuries

We examined female mice considering the relatively understudied PT-OA in females despite a higher risk of PT-OA^50, 51^ and ACL-tear in females than males^14, 52-54^. PT-OA was induced by a non-invasive ACL-injury^55^ in 8-10-week-old female C57BL/6 mice unilaterally (injury in one knee) or bilaterally (injuries in both knees). Briefly, mice were anesthetized and placed on a custom-built Styrofoam. A custom-built strain gauge-instrumented probe (Fig. 1a-b) was placed onto the skin over the patellar tendon, and an axial force was manually applied to the distal femur along the axis of the femoral shaft. The temporal force profiles were recorded using an Arduino system. The average rupture force was 14.2±0.3 N and the loading rate was 1.52±0.05 N/s (n=94 limbs) (Fig. 1c-e). Mice had positive Lachman tests and X-ray imaged were acquired immediately after the procedure to ensure that the injury had not caused a fracture of the tibia or femur. Mice were monitored for 3 days for signs of pain. This procedure was approved by the University of Rochester Committee on Animal Resources (UCAR). (see supplemental material for the detailed method).

**Fig 1.**
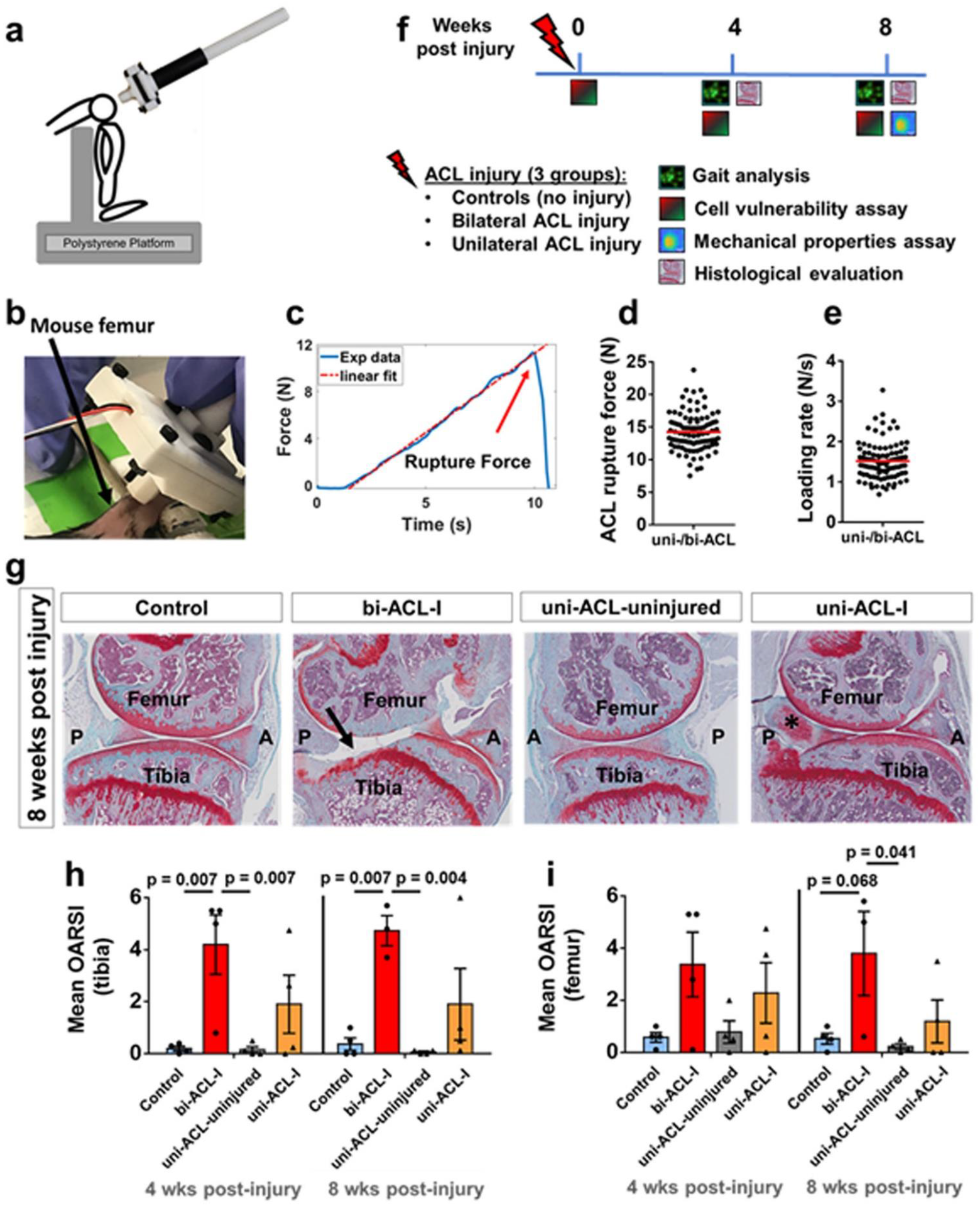
N*on-invasive ACL-injury method and study design*. (**a**) Schematic representation and (**b**) a representative figure of the experimental setup where the ACL injuries were induced with a custom-built strain gauge-instrumented device by applying axial force along the femoral shaft until a distinct sound of ACL tear was heard and the force drop was felt. The method was adapted from [Zhang 2020]. (**c**) Representative force-time curve obtained during the ACL injury procedure. The rupture force is indicated by the arrow, and the dashed line indicates linear curve fit from which the loading rate was calculated. (**d, e**) Quantification of ACL rupture force and loading rate. Each data point represents ACL injured limb and red line denotes the mean of the data. (**f**) Study design. ACL injuries were performed on either one knee joint (uni-ACL-I group), or on both knee joints (bi-ACL-I group) in 8-9-week-old female C57BL/6 mice. Gait, cell vulnerability, mechanical properties, and histological analysis were assessed at different timepoints post ACL injury. (**g-i**) Histological evaluation of articular cartilage in murine knee joints in 3 experimental groups (Control, bi-ACL-I, uni-ACL: injured and contralateral uninjured) at 4-, and 8-weeks post ACL-I timepoints. (**g**) Representative SaFO/FG-stained sagittal histological sections of medial side of mouse knee joints in control, bi-ACL-I, uni-ACL-I, and the contralateral uni-ACL-uninjured 8 weeks post ACL injury. Significant articular cartilage degradation (arrow) is evident on the bi-ACL injured knee, indicative of PTOA. Tissue calcification (*) was also observed on the uni-ACL-injured knees. A and P denote anterior and posterior direction of a knee joint. **(h, i)** Semiquantitative assessment of cartilage degradation on mouse (**h**) proximal tibias and (**i**) distal femurs using OARSI scoring system (n = 3-4 mice/group). Each data point represents an average of blinded scoring by 4 individuals. Higher OARSI scores (max = 6) indicate a higher degree of cartilage degradation. OARSI scores were compared using a Two-way ANOVA with a post-hoc Tukey test. Data are mean ± SEM. p≤0.05 indicates statistical difference between the groups.

### Gait analysis

Alteration in mouse gait post-ACL-I was assessed by Noldus CatWalk XT automated gait analysis system (Noldus Information Technology, XT10.6)^56, 57^. Briefly, a mouse walked across an enclosed space on top of a 50 cm long glass plate illuminated by an internally reflected green LED light. Below the glass, a high-speed color camera captured green light refraction of the illuminated mouse paw prints when the paw touched the glass. Red light illuminated the top of the glass plate to capture mouse silhouette during the gait (Fig. 2a-b). Each mouse walked 3 compliant runs with a run variance below 65%. Gait analysis was performed longitudinally at 4- and 8-weeks post-injury (n = 6/group). The hindlimb maximum Contact Mean Intensity was measured from the captured timeframe with the most intense pixels of the paw prints during a mouse’s gait cycle indicating max mouse weight-bearing. The max Contact Mean Intensity of left and right hind limbs in bi-ACL-I group were averaged and compared to other groups. The Base of Support, a vertical distance between the hind limbs during the gait, was also measured to quantify mouse posture (Fig. 2e-f). The Stand mean, a contact duration of hind paws on the walkway glass, was compared between groups (Suppl. Fig. 1). There was no difference in average mouse weight or running speed between uni-ACL-I and bi-ACL-I mice (Fig. 2d, g).

**Fig 2.**
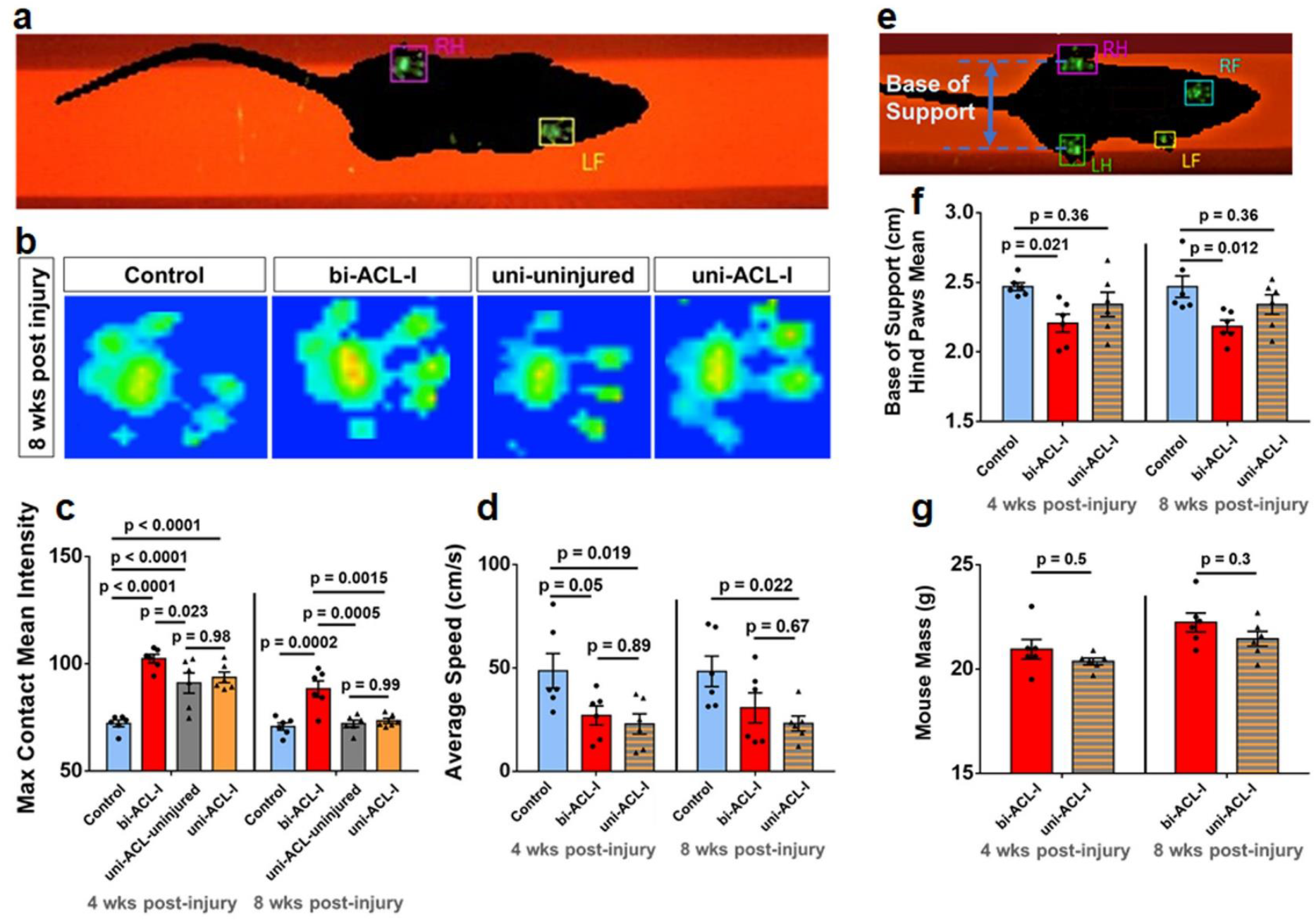
Bi-ACL-I and uni-ACL-I exhibit distinct weight bearing during locomotion. (**a)** Representative snapshot of intensity of mouse paw-prints on illuminated platform of a Noldus CatWalk XT system during gait analysis. (**b**)Representative snapshots of hind paw-print intensities at 8-weeks post-injury in different experimental groups. See supplemental videos of representative 3D contact intensity plots during mouse locomotion. (**c**) Quantification of maximum contact mean intensity in experimental and control groups. Note that in bi-ACL-I and uninjured control mice max contact mean intensities were averaged for both hind limbs per mouse, while uni-ACL-I and uni-ACL-uninjured hindlimbs were treated separately. (**d**) Average mouse speed in 3 experimental groups (Control, bi-ACL-I, uni-ACL-I) at 4- and 8-weeks post-ACL-I. (**e**) Representative micrograph of a mouse on a CatWalk platform with the indicated base of support as the width between mouse hindlimbs during locomotion, and (**f**) quantification of mean base of support. (**g**) Measurement of mouse mass in bi-ACL-I and uni-ACL-I groups prior to gait analysis at 4- and 8-weeks post-injury. 6 mice per group were used to longitudinally assess their gait. Contact intensities, average speeds, base of support, and mouse mass were compared using repeated measures Two-way ANOVA with a post-hoc Tukey test. Data are mean ± SEM. p≤0.05 indicates statistical difference between the groups.

### Mechano-vulnerability assay (Impact-induced chondrocyte LIVE/DEAD assay)

We quantified the vulnerability of chondrocytes on the *lateral* femoral condyles by 1 mJ impact^58^. Briefly, distal femurs were dissected and stained with calcein-AM and Propidium Iodide (PI) (R37601, Invitrogen), positioned in our custom-built impact device (Fig. 3a), then subjected to a 1 mJ kinetic energy on the patello-femoral groove. Chondrocytes on the lateral femoral condyles were z-stack imaged by a confocal microscope (FV3000, Olympus, UPlanSApo 10X/0.40 NA dry) before and after the impact. After the impact, specimens were re-stained with PI, an indicator of injured/dead cells, and re-imaged. Chondrocytes that lost calcein-labeling and became PI-positive were considered to be injured/dead, and the area of injured cells was quantified using ImageJ. (see supplemental material for the detailed method).

**Fig. 3.**
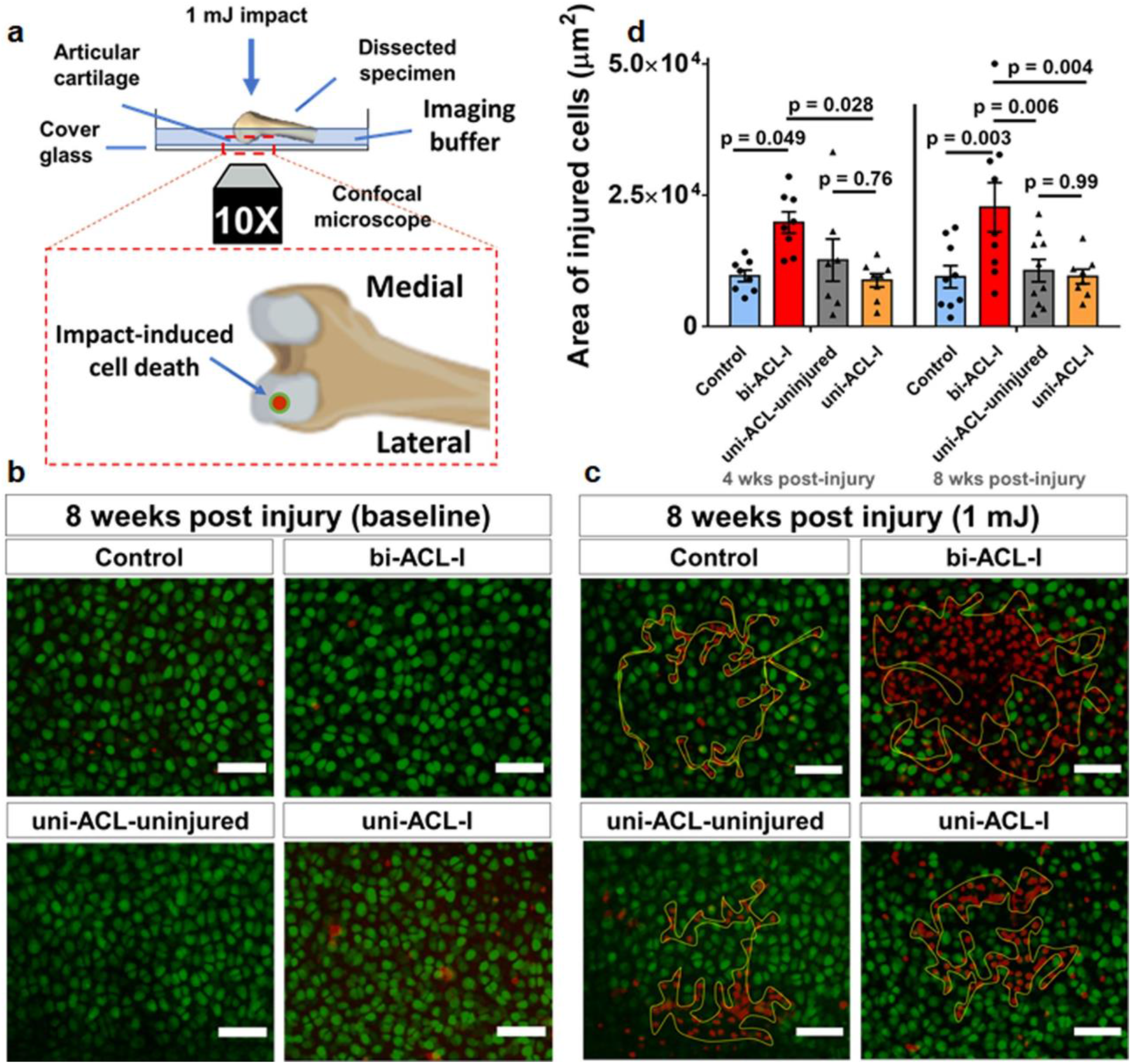
Chondrocytes in bi-ACL-I mice are more vulnerable than in uni-ACL-I mice. (**a**) Schematic representation of mechano-vulnerability experimental setup. Vitally stained *in situ* chondrocytes on lateral condyles of harvested distal femurs were imaged before and after the application of 1 mJ kinetic energy impacts. A zoomed-in inset shows a view of femoral condyles in the imaging plane where an impact-induced cell death is analyzed on the lateral condyles. (**b, c**) Representative z-projections of confocal micrographs of live/dead (green/red) chondrocytes acquired (**b**) before and (**c**) after the impact at 8 weeks post injury. Yellow contours in (**c**) indicate areas of injured/dead cells due to the applied 1mJ impact. The scale bars are 50 mm. (**d**) Quantification of impact-induced areas (within yellow contours) of injured cells on surface of lateral femoral condyles in the experimental groups at 4- and 8-weeks post ACL-I time-points (n = 6-9 limbs/group). The areas were compared using a Two-way ANOVA with a post-hoc Tukey test. Data are mean ± SEM. p≤0.05 indicates statistical difference between the groups.

### Mechanical properties of articular cartilage

A confocal microscope-based inverse finite element method (iFEM)^59^ was used to quantify the altered intrinsic stiffness (Young’s Modulus) of the ECM at 8-weeks post-injury. In brief, distal femurs were stained with 5’-DTAF (Sigma-Aldrich) to label the cartilage ECM, and placed on a cover glass of a custom-built mechanical device above a confocal microscope (Olympus FV3000, LUCPLFLN 40X NA = 0.6) (Fig. 2a). Confocal z-stacks of lateral femoral condyles were obtained before (baseline) and 5 min after applying a static load of 0.1 N (0.31 μm/pixel in xy-plane and 0.89 μm/slice in z-plane). The acquired confocal z-stacks were used to obtain thickness ratios (i.e., compressive tissue stretch λ_*z*_), maps of the infinitesimal tissue strain (ε_*z*_ = 1 − *thickness ratio*), and peak compressive strains (strains within 5 μm from the peak compressive strain). The compression experiments were simulated in FEBio using 3D FEMs^60^ to determine the Young’s modulus of the femoral ECM in different experimental groups (see supplemental material for detailed methods).

### Histological evaluation

After 4- and 8-weeks post-injury, mouse tibio-femoral joints were fixed in 10% formalin, decalcified, and embedded in paraffin. Sagittal sections (7 μm thick) of medial knee joints were stained with Safranin-O/Fast Green (Saf-O/FG, Applied Biosciences), and imaged by Virtual Slide Microscope System (Olympus VS120, 40X dry). Osteoarthritis Histopathology Assessment System (OARSI) scores^61^ of femur and tibia cartilage are assessed in a blind-manner.

### Statistical analyses

Cell death areas and OARSI score) were compared between the experimental groups using a Two-way Analysis of Variance (ANOVA) with a post-hoc Tukey test. Max Contact Mean Intensity, Base of Support and mouse mass were compared across experimental groups using repeated measures Two-way ANOVA with a post-hoc Tukey test. Compressive strains and Young’s moduli of ECM were compared using a One-way ANOVA with a post-hoc Tukey test. Statistical differences were detected at a significance level (α) of 0.05.

## Results

To investigate the role of *in vivo* loading in PT-OA development post-ACL-I, we compared gait parameters of hind limbs, mechano-vulnerability of articular chondrocytes *in situ*, mechanical properties of the ECM, and OARSI score in experimental groups: mice without injury (Group 1: uninjured controls), mice with bilateral ACL injury (Group 2: bi-ACL-I), and mice with a unilateral ACL injury, where the contralateral knee joints served as the unilateral uninjured controls (Group 3: uni-ACL-uninjured, Group 4: uni-ACL-I) at 0∼8 weeks post-injury (Fig. 1f-i).

### Bilaterally ACL-injured mice experience higher weight-bearing in their hindlimbs during the gait than unilaterally ACL-injured mice

We first found that the bi-ACL-I joints exhibit significantly increased hindlimb Max Contact Mean Intensity (related to mouse weight-bearing, arbitrary unit (a.u.)) during mouse locomotion as compared to uni-ACL-I mice and uninjured control group at both 4- and 8-week post-injury (Fig. 2a-b; week 4: 102.4±1.98 a.u. vs. 72.3±1.54 a.u., week 8: 88.3±3.72 a.u. vs. 70.8±1.67 a.u., mean±SEM). In contrast, uni-ACL-I joints exhibit an increased Contact Intensity as compared to the uninjured control group only at 4-week (93.7 ± 2.45 a.u.) but not at 8-week post-injury (73.5±1.16 a.u.). Interestingly, contralateral uni-ACL-uninjured joints exhibit similar levels of joint loading at both 4- and 8-week post-injury (Fig. 2a-b). Next, there were noticeable alterations in mouse posture during the gait in terms of the Base of Support, the vertical distance between mouse hind limbs. Bi-ACL injured mice had a narrower Base of Support as compared to uninjured control mice, while mice in the uni-ACL group had a Base of Support similar to the uninjured controls (Fig. 2c). The average duration of the mouse paw print (stand mean) was similar across all experimental groups and time-points (Supplemental materials). We note that the walking speed and bodyweight of bi-ACL-I and uni-ACL-I mice were similar for the entire course of the experiment (4- and 8-weeks post-injury) allowing for fair comparisons of the analyzed gait parameters; yet these uni-ACL- and bi-ACL-injured mice walked slower than uninjured control mice. These differential data of Contact Mean intensity and Base of Support reveal that the injured cartilage in bi-ACL-I joints experiences higher *in vivo* joint-loading and more significant mechanical instability during gait than injured cartilage in uni-ACL-I.

### *In situ* chondrocytes in bi-ACL-I joints are more vulnerable to impact loading than chondrocytes in uni-ACL-I or uninjured joints

Chondrocyte survival is implicated in the pathogenesis of PT-OA, and cell viability and metabolism under physiologic and pathophysiologic mechanical environments is crucial for cartilage remodeling and homeostasis. Therefore, we quantified areas of chondrocyte death in load-bearing cartilage on femoral condyles induced by application of injurious 1mJ impacts onto isolated cartilage-on-bone explants (Fig. 3a)^58^. This assay examines the mechano-vulnerability of *in situ* chondrocytes in load-bearing femoral cartilage, and the quantified areas of dead cells are presumed to represent the resilience of load-bearing chondrocytes against injuries *in vivo*. Interestingly, we found that bi-ACL-I joints exhibit the largest areas of cell death induced by impact loading at both 4- and 8-weeks post-injury. This result indicates that chondrocytes in bi-ACL-I became significantly more mechano-vulnerable than chondrocytes in uni-ACL-I or uninjured cartilage (Fig. 3b-c). In contrast, areas of dead cells in femurs of uni-ACL-I were comparable to contralateral uninjured joints (uni-ACL-uninjured) and uninjured controls (control group) at both 4- and 8-weeks post-injury (Fig. 3b-c). This result reveals that chondrocytes in uni-ACL-I maintained their mechano-vulnerability as uninjured chondrocytes. We also observed that, at 0-week (3-5 days) post-injury, areas of chondrocyte death were significantly elevated in both the uni- and bi-ACL-I groups as compared to uninjured controls. This significantly elevated mechano-vulnerability at 0-week is presumed to be driven by acute inflammation post-injury in synovial joints^62, 63^ (Control: 11707.8 ± 1081.4 μm^2^, bi-ACL-I : 16312.1 ± 901.1 μm^2^, uni-ACL-uninjured: 13449.8 ± 535.3 μm^2^, uni-ACL-I: 22106.4 ± 679.7 μm^2^; Supplemental material). Taken together, these data reveal that articular chondrocytes in bi-ACL-I joints become more vulnerable from injurious mechanical loading as compared to the chondrocytes in uni-ACL-I or uninjured joints as developing PTOA (Fig. 3).

### Mechanical property of ECM: soften in uni-ACL-I, stiffen in bi-ACL-I at 8-weeks post-injury

We measured alterations in ECM Young’s modulus and cartilage thickness of lateral condyles at 8-weeks post-injury (Fig. 4, Supplemental material). We found that articular cartilage of uni-ACL-I had reduced Young’s moduli and increased peak compressive infinitesimal strains relative to contralateral uninjured limbs (3.6±0.3 MPa vs. 7.8±1.5 MPa; 35.4±1.1 % vs. 26.6±2.5 %; Fig. 4d-e), indicating cartilage softening and the onset of PTOA only in the injured limb but not in intact contralateral limb post-ACL-I. Interestingly, Young’s modulus of cartilage in bi-ACL-I joints was almost two-fold higher than that of uninjured controls (11.4±1.2 MPa vs. 6.1±0.7 MPa; Fig. 4e). Cartilage thickness of bi-ACL-I was significantly reduced as compared to uninjured controls (29.8±0.8 μm vs. 34.5±1.1 μm; Supplemental material), suggesting cartilage degradation and erosion. Though not significant, reduction in cartilage thickness was also observed in both uni-ACL injured and uni-ACL-uninjured limbs as compared to uninjured controls (Supplemental material). Taken together, ECM of articular cartilage in uni-ACL-I group was less stiff and the most compressed by a static load among the groups, whereas bi-ACL-I group exhibited the stiffest and least compressed cartilage. These differences in mechanical properties of the ECM suggest that *in vivo* chondrocytes may experience distinct mechanical stimuli post-injury in our experimental groups (bi-ACL-I, uni-ACL, and uninjured controls), which lead to a feed-forward distinct cellular mechano-sensitivity and biosynthetic activities, which in turn alter structural and mechanical properties of the ECM along the PT-OA development.

**Fig. 4.**
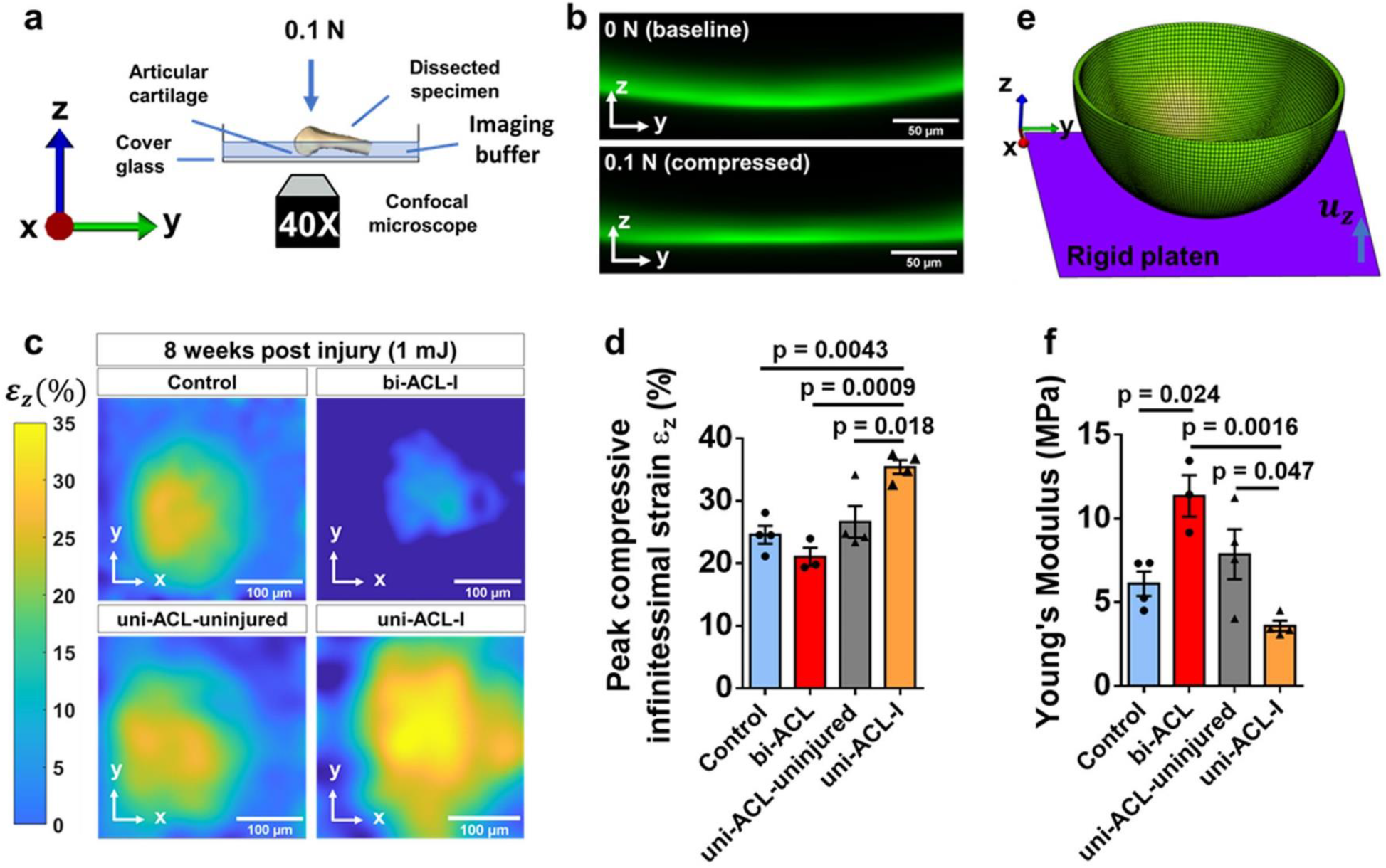
The ECM of Femoral lateral condyles in bi-ACL-I hindlimbs are stiffer than the ECM in uni-ACL-I hind limbs. (a) Schematic representation of experimental setup to assess compressive strain and mechanical properties of cartilage ECM on lateral femoral condyles. (b) Representative confocal micrographs of fluorescently stained cartilage on the lateral femoral condyle (top) before and (bottom) during cartilage compression after the specimens were subjected to a 0.1 N static loading for 5 min. Scale bar is 50 mm. (c) Representative maps of compressive strain of cartilage ECM on lateral femoral condyles of dissected specimens subjected to a 0.1 N static loading for 5 min in 3 experimental groups (Control, bi-ACL-I, uni-ACL: injured and contralateral uninjured) at 8-weeks post ACL-I time-point (n = 3-4 mice/group). Scale bar is 100 mm. (d) Quantification of the peak compressive infinitesimal strain in the experimental groups. (e) Representative 3D geometry of cartilage FEM to simulate the compression experiment and quantify ECM Young’s modulus via an inverse FE analysis. The cartilage is compressed with a rigid plated with prescribed experimentally measured boundary displacements (u_z_). (f) Quantification of solid matrix Young’s modulus of cartilage on lateral femoral condyles in the experimental groups using a finite element-based method and parameters including reaction force, boundary displacement, and cartilage geometry (see Supplementary materials). Compressive strains and Young’s moduli were compared using a One-way ANOVA with a post-hoc Tukey test. Data are mean ± SEM. p≤0.05 indicates statistical difference between the groups.

### Bi-ACL-I joints develop more severe PT-OA than uni-ACL-I joints

Histological sections of the medial knee joints revealed that uni-ACL-I and bi-ACL-I induce different degrees of cartilage degeneration at 4- and 8-weeks post-injury (Fig.1f-g). Specifically, in the majority of bi-ACL-I joints, we observed an anterior shift of mouse tibias and menisci relative to the femurs. In addition, significant articular cartilage degradation occurred on the posterior side of the tibia in bi-ACL-I. In contrast, the majority of uni-ACL-I knees did not have meniscus shift, cartilage degradation on the posterior side of the tibia was less severe than in bi-ACL-I group, and tissue calcification was observed (Fig. 1g, * marked region). The mean OARSI scores of cartilage on medial femoral condyles were 0.6 and 0.5 in uninjured control, 3.4 and 3.8 in bi-ACL-I, 2.3 and 1.2 in uni-ACL-I, and 0.6 and 0.5 in uni-ACL-uninjured at 4- and 8- weeks post-injury, respectively (Fig. 1h-i). The mean scores within the medial tibial plateau (MTP) were 0.2 and 0.4 in uninjured controls, 4.2 and 4.7 in bi-ACL-I, 1.9 and 1.9 in uni-ACL-I, and 0.2 and 0.1 in uni-ACL-uninjured at 4- and 8-weeks post-injury, respectively (Fig.1h-i). These results suggest that mice with bi-ACL-I develop more severe or rapid PT-OA as compared to uni-ACL injured mice.

## Discussion

Joint injury is a risk factor for OA, and altered joint kinematics due to joint injury has been suggested to contribute to PT-OA development, whether by directly damaging chondrocytes and ECM or indirectly altering chondrocyte metabolism and ECM stiffness progressively. Understanding of cartilage mechanotransduction may shed the light into developing effective therapeutic strategies to treat PT-OA^6, 64-68^. Here, we made a step towards understanding the relationship between the *in vivo* joint loading and chondrocyte mechano-vulnerability in murine articular cartilage post unilateral or bilateral ACL-I. Our data revealed that a higher *in vivo* joint loading leads to more severe or more rapid PT-OA development post-ACL-I. Interestingly, articular chondrocytes in the more loaded femoral cartilage (i.e., in bi-ACL-I group) were more mechano-vulnerable and more prone to cellular death by mechanical stimuli as compared to chondrocytes in the less loaded cartilage (i.e., in uni-ACL-I or uninjured control groups). Our findings suggest that it is critical to avoid abnormal joint mechanics and to tightly tune the chondrocyte mechanosensitivity post-ACL-I in order to slow down PT-OA progression.

Unilateral and bilateral ACL-I differentially altered mouse gait patterns. We found that weight-bearing in hind limbs of bi-ACL-I mice was higher than in uni-ACL-I limbs and in uninjured control limbs, while the walking speed and body weight were comparable between bi-ACL-I and uni-ACL-I mice at both 4- and 8-weeks post-injury. Since uni-ACL-I destabilizes one hindlimb knee joint and the contralateral joint stays intact, we anticipated that uni-ACL-I mice exhibit asymmetrical gait distribution with lower load-bearing on injured limbs. Contrary to our expectations and consistent with previous reports indicating that mice can restore a symmetric distribution of weight-bearing within 2-3 weeks following traumatic injury^62, 69^, uni-ACL-I mice compensated the unbalanced joints, and exhibited similar levels of weight-bearing in injured and contralateral uninjured hind limbs at 4- and 8- weeks post-injury. Another distinctive difference between uni-ACL-I and bi-ACL-I mice was in the Base of Support, the width of hind-limb positions during the mouse locomotion. We observed a significantly reduced Base of Support in bi-ACL-I mice at 4-, and 8-weeks post-injury, while uni-ACL-I mice showed a comparable length of the Base of Support with uninjured control mice. These data indicate that unilateral and bilateral ACL-I cause differential mechanical environments in injured limbs that can contribute to different degrees of PT-OA development, and allows to examine the effect of joint mechanics on cellular and tissue health.

Distinct cellular mechano-vulnerability was observed in articular chondrocytes in lateral femoral condyles of uninjured, uni-ACL-I and bi-ACL-I joints over PT-OA development. Since chondrocyte death precedes cartilage degeneration in OA, and thus by promoting chondrocyte survival post-injury would delay PT-OA pathogenesis, we examined the *in situ* chondrocyte survival due to the applied mechanical impacts post-ACL-I. At 0-week post-injury, we observed increased areas of chondrocyte death induced by 1mJ mechanical impacts in ACL-I joints (uni- and bi-ACL-I) as compared to the areas observed in uninjured joints (Supplemental material). This increased mechano-vulnerability of chondrocytes in uni-ACL-I and bi-ACL-I groups is presumably due to the local acute inflammation as the early response of synovial joints to trauma^62, 70^. Interestingly, at both 4- and 8-weeks post-injury, articular chondrocytes in femoral condyles of bi-ACLI-I joints were the most vulnerable to mechanical impacts than chondrocytes in the uninjured control group (Fig. 3). The vulnerability of articular chondrocytes in uni-ACL-I joints were comparable to the chondrocytes’ vulnerability in contralateral uninjured joints or to uninjured control limbs at 4- and 8-week post-injury. Importantly, these trends in cell mechano-vulnerability are similar to the trends of hindlimb weight bearing observed during mouse locomotion. We also observed that baseline cell death (cell death before the application of mechanical impacts) was elevated on medial femoral condyles as compared to cell death on lateral condyles, which is consistent with a study conducted by Berke et al.^62^. Therefore, we anticipate that medial cartilage endures higher degrees of abnormal mechanical loads post-ACL-I than lateral cartilage does, thereby causing more rapid cartilage degradation by altering chondrocyte mechano-sensitivity and cartilage integrity.

Distinct mechanical properties of cartilage ECM were also observed in lateral femoral condyles of uni-ACL-I and bi-ACL-I joints at 8-weeks post-injury. Based on the equilibrium Young’s moduli, uni-ACL-I limbs have significantly lower ECM modulus as compared to ECM in contralateral uninjured or control uninjured limbs at 8-weeks post-injury. This reduction in ECM modulus corresponded to moderate PT-OA levels with 2∼4 OARSI scores (Fig. 1g-i). However, and to our surprise, bi-ACL-I knee joints showed a significantly increased Young’s moduli of femoral articular cartilage and the most severe OARSI scores than other experimental groups (Fig. 1g-i). Altered cartilage moduli indicate OA progression and cartilage erosion in PT-OA^71-74^. Doyran et al. has reported a similar transition of cartilage stiffness over PT-OA development in a murine destabilized medial meniscus (DMM) model^71^. In their animal model, the ECM of medial condyles of injured limbs had decreased nanoindentation modulus as compared to the ECM of sham-surgery limbs at 1∼8-weeks post-DMM-injury, then the modulus recovered significantly at 12-weeks post-injury when OA was the most severe based on their histological analysis^71^. Articular cartilage has a depth-dependent gradient in its compressive modulus with the less stiff superficial cartilage layer and the stiffer cartilage in deeper layers reaching the bone^75, 76^. Cartilage thickness of femoral condyles was reduced in both uni-ACL-I and bi-ACL-I, yet bi-ACL-I showed slightly more reduced cartilage thickness and higher radius of curvature than uni-ACL-I. We speculate that the stiffening of the ECM in bi-ACL-I was due to the more severe degradation of the superficial cartilage layer than in uni-ACL-I joints. Taking together, our ECM modulus and histology data indicate that bi-ACL-I joints have more severe OA than uni-ACL-I joints, and cartilage of bi-ACL-I is stiffer than cartilage of uni-ACL-I. Ironically, despite stiffened ECM, articular chondrocytes in bi-ACL-I were more sensitive to mechanical impacts at 4- and 8-weeks post-injury. Stiffened ECM did not lessen the effect of mechanical impacts on *in situ* chondrocytes, but increased impact-induced chondrocyte death in bi-ACL-I joints. These data suggest that chondrocytes in bi-ACL-I were more fragile and impaired more upon mechanical stimuli.

Considering the crucial role of chondrocyte viability in cartilage homeostasis, our results suggest that the increased mechano-vulnerability of chondrocytes contributes to an accelerated PT-OA pathogenesis. Thus, we anticipate the homeostasis of cellular mechano-sensitivity, or mechanical-injury-sensitivity, is critical to delay PT-OA. A group of mechanosensing and mechanotransducing molecules may be regulated over PT-OA development and tune cellular mechano-sensitivity under abnormal joint mechanics and inflammation post-ACL-I. For instance, dysregulated integrin αVβ_3_ and their associated ligands may play essential roles in disrupting chondrocyte-ECM interactions over OA progression^77-79^, as well as Piezo1 may be augmented via IL-1-mediated inflammatory patho-mechanisms^80, 81^. Future research will investigate the chondrocyte mechano-vulnerability based on mechanosensors and mechanotransducers and their local ECM properties over PT-OA development.

## Conclusion

In this study, we quantified chondrocyte vulnerability and ECM mechanics on articular cartilage in load-bearing knee femoral joints over OA development post ACL-injury. We found the *in vivo* joint loading-dependent changes on chondrocytes’ mechano-vulnerability over the PT-OA development in female mice post-ACL-I. To our knowledge, this is the first study comparing the mechano-vulnerability of chondrocytes and the mechanical properties of ECM between unilateral and bilateral ACL-injuries in female mice. Our data reveal that unilateral ACL-I mice compensate and balance their joint loading between injured and uninjured hind limbs, resulting in the delayed progression of PT-OA with minimal changes on cellular mechano-vulnerability as compared to injured knees of bilateral ACL-I mice. This study suggests that the reduction in knee joint loading may delay ACL injury-associated progression of OA. Furthermore, patients with injuries in both of their joints may have a higher risk to develop PT-OA than the patients who do not have another joint injury in their contralateral knees because of the increased joint-loading. Our results support therapeutic interventions to tune cellular mechano-sensitivity and physical therapy to correct aberrant joint loading in ACL-injured legs to slow down PT-OA progression.

## Additional materials and methods

Additional materials and methods are provided in Supplementary files.

## Author contributions

WL & SM conceived and designed the study. AK, AE, NYC and AP performed experiments. AK, AE, NYC, CP and WL analyzed and interpreted the data. AK, AE, NYC, SM, CP and WL were involved in drafting the manuscript for important intellectual content and all authors approved the final version of the manuscript.

## Acknowledgements

The authors thank Dr. Robert Dirksen for continuous research discussion. We also thank Center for Musculoskeletal Research (CMSR) Histology Core and University of Rochester Committee on Animal Resources (UCAR) for their technical assistance.

