## Supplementary material for "Effect of knee joint loading on chondrocyte mechano-vulnerability and severity of post-traumatic osteoarthritis induced by ACL-injury in mice": Suppl Material

### 1. Supplementary Materials and Methods

#### 1.1. Mechano-vulnerability assay (Impact-induced chondrocyte LIVE/DEAD assay)

We quantified the vulnerability of chondrocytes in cartilage on the *lateral* femoral condyles by 1 mJ impact post ACL-I [58]. Briefly, immediately following euthanasia, distal femurs were dissected and stained with 10 μM calcein-AM and 60 μM Propidium Iodide (PI) (R37601, Invitrogen). The dissected specimens were positioned to face the cartilage on medial and lateral femoral condyles towards the glass-bottom of our custom-built impact device (Fig. 3a), then subjected to a 1 mJ kinetic energy impact on the patello-femoral groove (the anterior side of the femur). Articular chondrocytes on the lateral femoral condyles were z-stack imaged by a confocal microscope (FV3000, Olympus) with an Olympus UPlanSApo 10X/0.40 NA dry objective before and after the application of 1 mJ impact. After the impact, the specimens were re-stained with PI, an indicator of injured/dead cells, and re-imaged. Chondrocytes that lost calcein fluorescence and became PI-positive were considered to be injured/dead, and the resulting area of injured cells on *lateral* condyles was quantified from z-projections of confocal z-stacks using ImageJ. Specimens were kept hydrated in imaging buffer containing 2 mM $CaCl_2$, 137 mM NaCl, 20 mM D(+)Glucose, 4.5 mM $MgCl_2$, 2.4 mM KCl, 10 mM HEPES and 35 mM Mannitol at pH 7.4 and 340 mOsm. In this assay, lateral femoral condyles were examined because of the relatively higher chondrocyte death on medial condyles of ACL-injured limbs prior to applying the impact. This observation is consistent with the published injury-induced murine PT-OA model [59].

#### 1.2. Mechanical properties of articular cartilage

A confocal microscope-based inverse finite element method (iFEM) [60] was used to quantify the altered intrinsic stiffness (i.e., Young's Modulus) of the ECM at 8-weeks post-ACL injury. In brief, dissected distal femurs were stained with 5'-DTAF (Sigma-Aldrich, 10 µg/ml, 1 h) to fluorescently label the cartilage ECM. As shown in Fig. 2a, specimens were placed on a cover glass to image distal femoral condyles facing the glass of a custom-built mechanical device above the lens of laser scanning confocal microscope (Olympus FV3000, LUCPLFLN 40X NA = 0.6). Confocal z-stacks of lateral femoral condyles were obtained before (baseline) and 5 min after cartilage was compressed with a static load of 0.1 N (0.31 µm/pixel in x-y plane and 0.89 µm/slice in z plane), which is sufficient for mechanical equilibrium [60]. The acquired 3D confocal z-stacks were used to obtain thickness ratios (i.e., compressive tissue stretch $\lambda_z$), maps of the infinitesimal tissue strain ($\varepsilon_z = 1 - thickness\ ratio$), and peak compressive strains (strains within 5 µm from the peak compressive strain), Spatially varying thickness ratios were calculated by fitting normalized depth-wise intensity profiles with Gaussian curves, and calculating the ratio of corresponding spatially varying fit parameters $\sigma$, proportional to the cartilage thickness ($thickness\ ratio = \frac{\sigma_{0.1\ N}}{\sigma_{0\ N}}$)[60]. This compression experiment was simulated in FEBio [61] using 3D finite element models (FEMs) to determine the Young's modulus of the ECM of cartilage on femoral lateral condyles in different experimental groups . The cartilage was approximated as a uniform hemispherical shell with a specimen-specific cartilage thickness (mean ± SEM: Control: 34.5 ± 1.05 µm, bi-ACL-I: 29.8 ± 0.7 µm, uni-ACL-uninjured: 31.2 ± 1.45 µm, uni-ACL-I: 30.7 ± 0.5 µm) and a radius of curvature (Control: 779.3 ± 22.68 µm, bi-ACL-I: 853.6 ± 31.9 µm ,uni-ACL-uninjured: 762.9 ± 4.4 µm , uni-ACL-I: 765 ± 24.9 µm) measured from confocal z-stacks at baseline (see supplemental material). The top/inner surface of the FEMs (representing bone-cartilage interface) was constrained from movement in all directions, and the bottom/outer surface (representing the articular surface) was allowed to deform. The boundary displacement ($u_z$), which was prescribed to a rigid platen at the cartilage-glass interface to deform the articular surface along the z-direction, was calculated from the experimentally measured peak infinitesimal strains ($u_z = \varepsilon_z^{peak} \times$ Cartilage thickness at the baseline). The cartilage was modelled as neo-Hookean hyper-elastic material with a Poisson ratio of 0.2 [62]. An iterative parameter optimization algorithm was used to determine Young's modulus of the cartilage ECM by matching the reaction forces observed on the rigid platen of FEMs to the forces measured experimentally (Supplemental material). Note that the 0.1 N force applied on top of the specimens during the experiment is distributed to three specimen-glass contacts (i.e., two cartilage-glass contacts on distal femoral condyles and one bone-glass contact on a proximal end of the explant). Cartilage-glass contact forces on lateral condyles of distal femurs were measured using a previously established method[60] (mean ± SEM, control: 0.04 ± 0.003 N, bi-ACL-I: 0.05 ± 0.002 N, uni-ACL-uninjured: 0.05 ± 0.0003 N , uni-ACL-I: 0.045 ± 0.001 N, see supplemental material). We note that this method quantifies bulk mechanical properties of the ECM with assumptions on cartilage structure and mechanical behavior. We simplified a complex poroelastic behavior of articular cartilage under compression with its known strain dependent permeability [63] and depth-dependent material properties [64, 65] , and instead, treated cartilage as homogeneous isotropic neo-Hookean elastic material, the quantification of which still allowed for adequate comparisons between experimental groups.

### 2. Supplementary Results

#### 2.1. Design of ACL-injury device

A custom-built strain gauge-instrumented device was designed in order to measure force applied during the ACL injuries over time. The device consists of a 3D-printed handle and hexagonal probe with cross-sectional area of 32.8 $mm^2$ printed using the polylactic acid material (Fig. S1a-b). An aluminum beam with a glued strain gauge was connected to the bottom of the handle and to the top of the probe, thereby serving as an interface between these two parts in the device (Fig. S1a-b). The axial force applied along the long axis of the handle during the ACL injuries (Fig. S1c) results in bending of the aluminum beam. The glued strain gauge, connected to an Arduino microcontroller through a load cell amplifier, outputs voltages that are linearly proportional to the applied force (i.e., the amount of beam bending) as indicated by a calibration curve (Fig. S1d). The knee joint dissections performed on some mice after the injury revealed an intact posterior cruciate ligament (Fig. S1e).

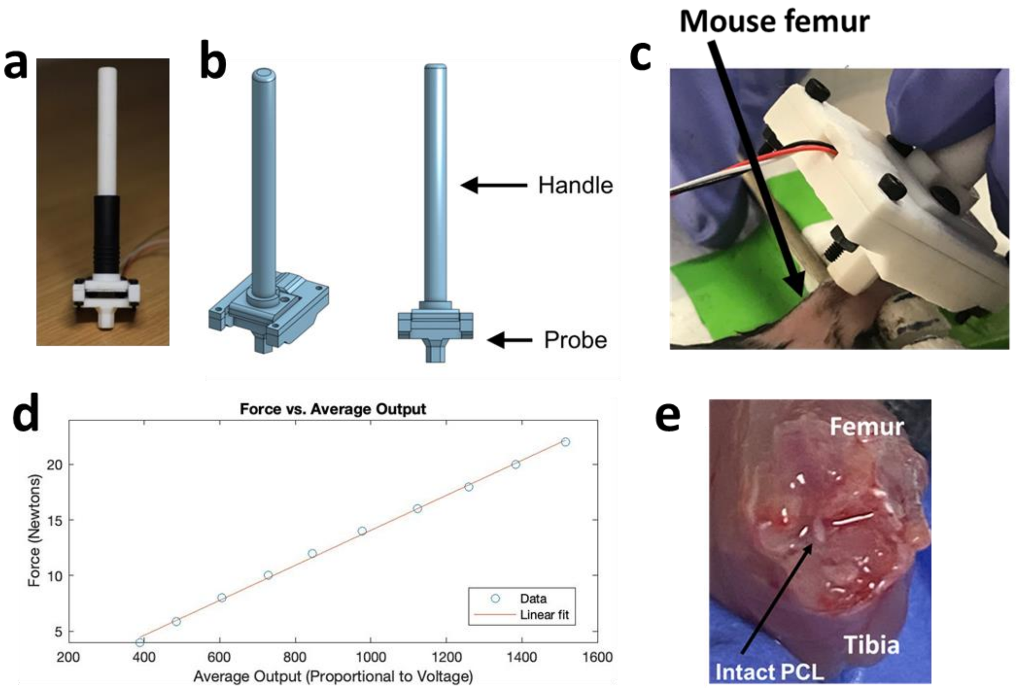


**Fig S1.** Visual representation of the device and non-invasive ACL injury. (a) Photograph of the ACL injury device. (b) CAD rendering of the device used for 3D printing. The ACL injuries were performed by (c) holding the handle and pressing the probe against mouse femur. (d) Calibration curve of digital output of the microcontroller vs. the applied force. 12 different forces ranging from 4 to 22 N were applied using an MTS machine, and the corresponding digital outputs from microcontroller were recorded over 15 seconds of the applied loads. A linear fit ($y = 0.0157x - 1.66, R^2 = 0.998$) was obtained in MATLAB. (e) Photograph of a dissected knee joint after the ACL injury showing an intact posterior cruciate ligament (PCL).

**2.2 Gait analysis: Quantification of stand mean**

Stand mean was quantified as the average duration of the hind paw contacting the walkway glass during the recorded trials. Stand means were averaged for the hind limbs (right and left limbs) per mouse in the control and bi-ACL-I groups, and separately quantified for injured and uninjured limbs in the uni-ACL group. All groups demonstrated similar levels of stand mean over the course of experiments (Fig. S2).

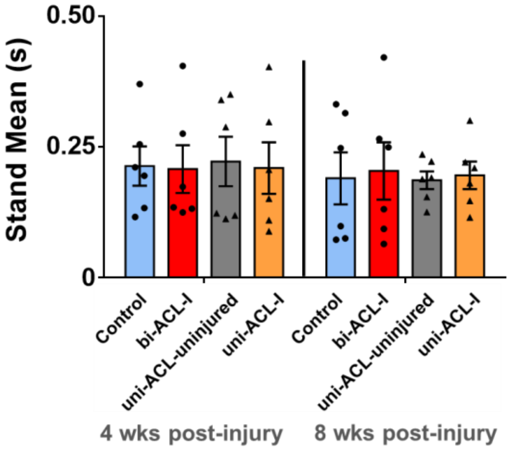


**Fig. S2**. *Quantification of stand mean during mouse locomotion (n = 6/group). The stand mean was compared using repeated measures Two-way ANOVA with a post-hoc Tukey test. Data are mean ± SEM.*

#### 2.3 Cell Vulnerability at 0 weeks post injury

An elevated cell vulnerability was observed in uni-ACL-I and bi-ACL-I experimental groups within 3 to 7 days post injury (0 weeks post injury timepoint, Fig. S3).

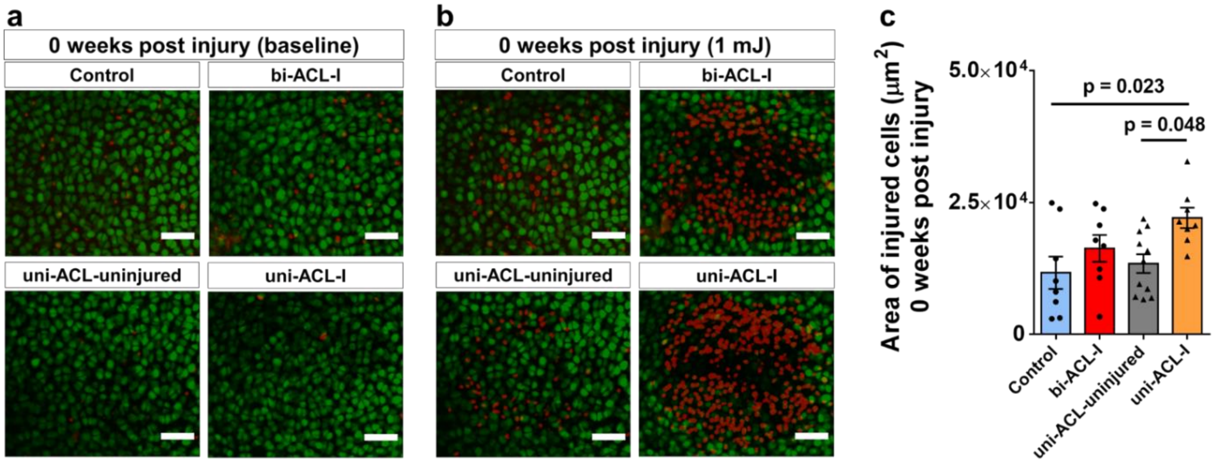


**Fig. S3.** ***(a, b)*** *Representative z-projections of confocal micrographs of live/dead (green/red) chondrocytes acquired* ***(a)*** *before and* ***(b)*** *after the impact at 0 weeks post- injury. Yellow con-tours in (b) indicate areas of injured/dead cells due to the applied 1mJ impact. The scale bars are 50 □m.* ***(c)*** *Quantification of impact-induced areas (within yellow contours) of injured cells on surface of lateral femoral condyles in the experimental groups at 0-weeks post ACL-I timepoint (n = 6-9 limbs/group). The areas were compared using a One-way ANOVA with a post-hoc Tukey test. Data are mean ± SEM. p≤0.05 indicates statistical difference between the groups.*

#### 2.4 Parameters used to assess ECM Young's Modulus.

The geometry of finite element models was determined from the experimentally measured cartilage thicknesses (Fig. S4a) and outer radii of curvature (Fig. S4b) obtained from confocal z-stacks of fluorescently labeled articular cartilage using a previously published method[1]. Briefly, sagittally reconstructed 2D images sliced at the center of lateral condyles (Fig. 4b) were used to measure cartilage thickness in ImageJ, and the outer radius of curvature in MATLAB by fitting a circle to the curvature of the articular surface. Though not statistically significant, a noticeable change in radius of curvature was observed in bilaterally injured mice (bi-ACL) at the 8-week timepoint post ACL injury (Fig. S4a). Cartilage thickness decreased in the 8-week post ACL injured groups as compared to uninjured controls with the statistical significance in the bi-ACL group (Fig. S4b), indicating cartilage degeneration during the progression of PTOA.

The contact (reaction) forces on lateral femoral condyles were experimentally assessed using a previously established method[1]. The applied 0.1 N force on top of the specimens creates three glass contact points: lateral condyle-glass, medial condyle-glass, and proximal bone-glass contacts. The contact force on lateral femoral condyles was quantified through a moment balance equation based on the location of the contact points and of the applied 0.1 N force. These contact locations were determined by placing specimens on a sheet covered with black ink and compressing them with a prescribed weight also covered with black ink. The specimens were then imaged under inverted and upright microscopes, and the contact locations were detected by the locations of the black prints on the specimens. To quantify the solid matrix Young's modulus of cartilage in different experimental groups, the specimen-specific reaction forces on the lateral condyles (Fig. S4c) were used to match the reaction forces observed in the FE simulations due to prescribed boundary displacements ($u_z$) (Fig. S4d). The boundary displacements were experimentally measured as the difference between thicknesses of before and during cartilage compression.

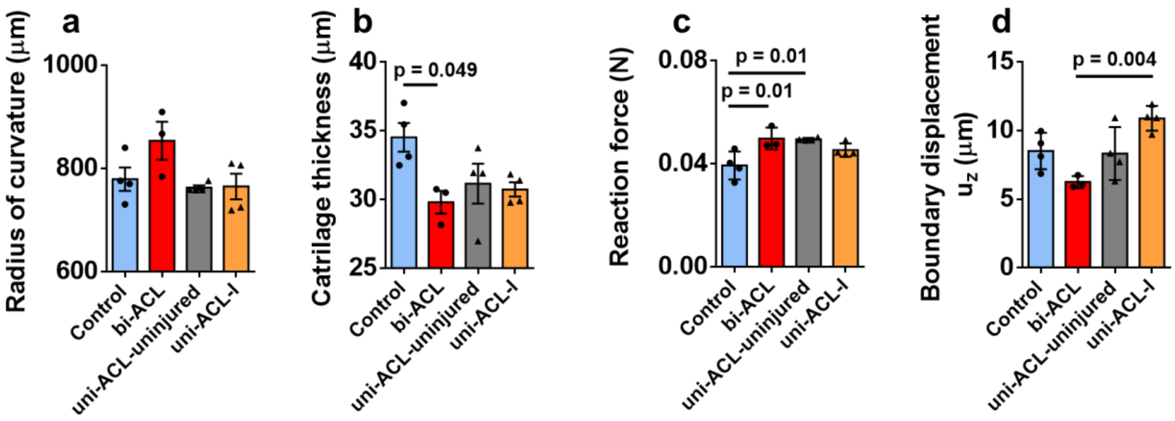


*Fig. S4 (**a**) Outer radius of curvature and (c) cartilage thickness were used to generate geometry of cartilage FEMs to simulate cartilage compression. (c) Reaction forces and (e) boundary displacements were used for inverse FEA to determine cartilage modulus. All the parameters (a-d) were compared between the experimental groups using a One-way ANOVA with a post-hoc Tukey test (n = 3-4 mice/group). Data are mean ± SEM. p≤0.05 indicates statistical difference between the groups.*
